# Gene model for the ortholog of *raptor* in *Drosophila erecta*

**DOI:** 10.64898/2026.07.09.737526

**Authors:** Anne E. Backlund, Jonas Nielsen, John Pulford, Brittany Cook, Jenna Anderson, Michelle Robert, Jeffrey S. Thompson, Chinmay P. Rele, Jacqueline K. Wittke-Thompson

**Affiliations:** The University of Alabama, Tuscaloosa, AL USA; Denison University, Granville, OH USA; University of St. Francis, Joliet, IL USA

## Abstract

Gene model for the ortholog of *raptor* in the May 2011 (Agencourt Dere_CAF1/DereCAF1) Genome Assembly (GenBank Accession: GCA_000005135.1) of *Drosophila erecta*. This ortholog was characterized as part of a developing dataset to study the evolution of the Insulin/insulin-like growth factor signaling pathway (IIS) across the genus *Drosophila* using the Genomics Education Partnership gene annotation protocol for Course-based Undergraduate Research Experiences.

## Introduction

*This article reports a predicted gene model generated by undergraduate work using a structured gene model annotation protocol defined by the Genomics Education Partnership (GEP; thegep.org) for Course-based Undergraduate Research Experience (CURE). The following information in quotes may be repeated in other articles submitted by participants using the same GEP CURE protocol for annotating Drosophila species orthologs of Drosophila melanogaster genes in the insulin signaling pathway*.

“In this GEP CURE protocol students use web-based tools to manually annotate genes in non-model *Drosophila* species based on orthology to genes in the well-annotated model organism fruit fly *Drosophila melanogaster*. The GEP uses web-based tools to allow undergraduates to participate in course-based research by generating manual annotations of genes in non-model species (Rele et al., 2023). Computational-based gene predictions in any organism are often improved by careful manual annotation and curation, allowing for more accurate analyses of gene and genome evolution (Mudge and Harrow 2016; Tello-Ruiz et al., 2019). These models of orthologous genes across species, such as the one presented here, then provide a reliable basis for further evolutionary genomic analyses when made available to the scientific community.” (Myers et al., 2024).

“The particular gene ortholog described here was characterized as part of a developing dataset to study the evolution of the Insulin/insulin-like growth factor signaling pathway (IIS) across the genus *Drosophila*. The Insulin/insulin-like growth factor signaling pathway (IIS) is a highly conserved signaling pathway in animals and is central to mediating organismal responses to nutrients (Hietakangas and Cohen 2009; Grewal 2009).” (Myers et al., 2024).

“raptor positively regulates (Target of Rapamycin) TOR-mediated cell apoptosis and growth control by differentially regulating S6K-dependent signaling pathways, and is a crucial regulator of cell growth and metabolism in *Drosophila* (Lee and Chung 2007; Hatfield et al., 2015). *raptor* is orthologous to the human *RPTOR* gene (*regulatory associated protein of raptor complex 1*) and is a well conserved component of the TORC1 complex (Wang et al., 2012).” (Backlund et al., 2025).

*“D. erecta* is part of the *melanogaster* species group within the subgenus *Sophophora* of the genus *Drosophila* (Sturtevant, 1939; Bock and Wheeler, 1972). It was first described by Tsacas and Lachaise (1974). *D. erecta* is found in west central Africa (https://www.taxodros.uzh.ch, accessed 1 June 2024; Markow and O’Grady, 2006) where it is found to breed primarily on the fruits of *Pandanus candelabrum*, a spiny evergreen shrub (Unwin, 1920; Lachaise and Tsacas, 1983).” (Lieser et al., 2024).

We propose a gene model for the *D. erecta* ortholog of the *D. melanogaster* raptor (*raptor*) gene. The genomic region of the ortholog corresponds to the uncharacterized protein LOC26526319 (RefSeq accession XP_015011450.1) in the Dere_CAF1 Genome Assembly of *D. erecta* (GenBank Accession: GCA_000005135.1 - Chirn et al., 2015; Ma et al., 2018). This model is based on RNA-Seq data from *D. erecta* (PRJNA414017, PRJNA264407) and *raptor* in *D. melanogaster* using FlyBase release FB2024_02 (GCA_000001215.4; Gramates et al., 2022; Jenkins et al., 2022; Larkin et al., 2021).

### Synteny

The target gene, *raptor*, occurs on chromosome X in *D. melanogaster* and is nested by *Multiple inositol polyphosphate phosphatase 2* (*Mipp2*). Furthermore, *raptor* is flanked upstream by *Ca2+-channel protein* α*1 subunit T* (*Ca-*α*1T*) and *Neprilysin 1* (*Nep1*) and downstream by *CG4660* and *CG4666*. The *tblastn* search of *D. melanogaster* raptor-PB (query) against the *D. erecta* (GenBank Accession: GCA_000005135.1) Genome Assembly (database) placed the putative ortholog of *raptor* within scaffold scaffold_4690 (CH954180.1) at locus LOC26526319 (XP_015011450.1)— with an E-value of 0.0 and a percent identity of 83.33%. Furthermore, the putative ortholog is nested within LOC6551318 (XP_015010628.1) and flanked upstream by LOC6551337 (XP_026837449.1) and LOC6551336 (XP_026837225.1), which correspond to *Mipp2, Ca-*α*1T*, and *Nep1* in *D. melanogaster* (E-value: 0.0, 0.0, and 0.0; identity: 93.86%, 95.05%, and 95.41%, respectively, as determined by *blastp*; Figure 1A, Altschul et al., 1990). The putative ortholog of *raptor* is flanked downstream by LOC6551316 (XP_026838083.1) and LOC6551315 (XP_026838084.1), which correspond to *CG4660* and *CG4666* in *D. melanogaster* (E-value: 0.0 and 9e-144; identity: 97.70% and 98.96%, respectively, as determined by *blastp*). The putative ortholog assignment for *raptor* in *D. erecta* is supported by the following evidence: The genes surrounding the *raptor* ortholog are orthologous to the genes at the same locus in *D. melanogaster* and synteny is completely conserved, supported by E-values and percent identities, so we conclude that LOC26526319 is the correct ortholog of *raptor* in *D. erecta* (Figure 1A).

**Figure 1.**
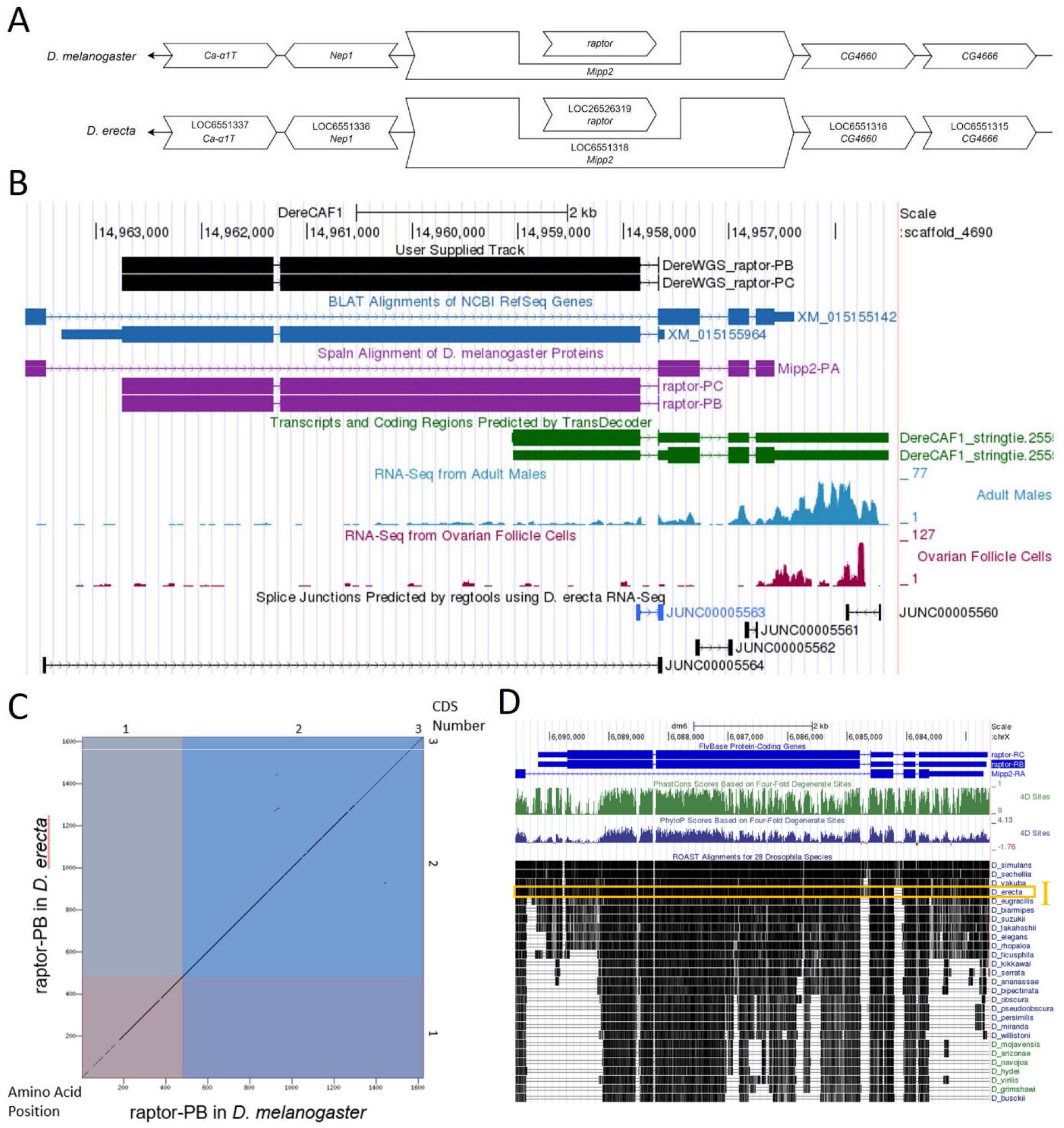
*raptor* gene model comparison between *Drosophila erecta* and *Drosophila melanogaster* orthologs. **(A) Synteny comparison of the genomic neighborhoods for *raptor* in *Drosophila melanogaster* and *D. erecta***. Thin underlying arrows indicate the DNA strand within which the target gene–*raptor*–is located in *D. melanogaster* (top) and *D. erecta* (bottom). The thin arrows pointing to the left indicate that *raptor* is on the negative (-) strand in both *D. erecta* and *D. melanogaster*. The wide gene arrows pointing in the same direction as *raptor* are on the same strand relative to the thin underlying arrows, while wide gene arrows pointing in the opposite direction of *raptor* are on the opposite strand relative to the thin underlying arrows. White gene arrows in *D. erecta* indicate orthology to the corresponding gene in *D. melanogaster*. Gene symbols given in the *D. erecta* gene arrows indicate the orthologous gene in *D. melanogaster*, while the locus identifiers are specific to *D. erecta*. **(B) Gene Model in GEP UCSC Track Data Hub (Raney et al., 2014)**. The coding-regions of *raptor* in *D. erecta* are displayed in the User Supplied Track (black); coding exons are depicted by thick rectangles and introns by thin lines with arrows indicating the direction of transcription. Subsequent evidence tracks include BLAT Alignments of NCBI RefSeq Genes (dark blue, alignment of Ref-Seq genes for *D. erecta*), Spaln of D. melanogaster Proteins (purple, alignment of Ref-Seq proteins from *D. melanogaster*), Transcripts and Coding Regions Predicted by TransDecoder (dark green), RNA-Seq from Adult Males and Ovarian Follicle Cells (light blue and dark red, respectively; alignment of Illumina RNA-Seq reads from *D. erecta*), and Splice Junctions Predicted by regtools using *D. erecta* RNA-Seq (PRJNA414017, PRJNA264407). Splice junctions shown have a read-depth of 1-10 and 10-49 supporting reads shown in black and blue, respectively. **(C) Dot Plot of raptor-PB in *D. melanogaster* (*x*-axis) vs. the orthologous peptide in *D. erecta* (*y*-axis)**. Amino acid number is indicated along the left and bottom; CDS number is indicated along the top and right, and exons are also highlighted with alternating colors. **(D) Conservation of 28 *Drosophila* species showing evolutionary conservation of raptor-PB**. Evidence tracks include FlyBase Protein-Coding Genes (blue), PhastCons Scores Based on Four-Fold Degenerate Sites (green), PhyloP Scores Based on Four-Fold Degenerate Sites (dark blue) and ROAST Alignments for 28 *Drosophila* Species indicating conservation relative to *D. melanogaster*. This region shows high local sequence similarity to *D. melanogaster* in multiple species. The ROAST alignment for *D. erecta* is highlighted by the gold box denoted I.

### Protein Model

*raptor* in *D. erecta* has two mRNA protein-coding isoforms (raptor-RB, raptor-RC; Figure 1B), containing three exons each. Relative to the ortholog in *D. melanogaster*, the CDS number and protein isoform count are conserved. The sequence of raptor-PB in *D. erecta* has 93.89% identity (E-value: 0.0) with the protein-coding isoform raptor-PB in *D. melanogaster*, as determined by *blastp* (Figure 1C). This model is supported by insufficient RNA-seq data in the coding regions of *raptor*. Additionally, a peak in RNA-seq data exists near the 3’ end of *raptor* (Figure 1B). This gene model can also be seen within the target genome at this TrackHub.

### Special characteristics of the protein model

#### Insufficient RNA-seq data in *D. erecta* to support the *raptor* gene model

The May 2011 (Agencourt Dere_CAF1/DereCAF1) assembly in *D. erecta* has Adult Male and Ovarian Follicle Cell RNA-seq data. The current RNA-seq data found in the *D. erecta* CAF1 assembly (Chirn et al., 2015; Ma et al., 2018) shows discontinuous and insufficient RNA-seq data within most of the coding sequence of *raptor* besides the 3’ end (Figure 1B). The peaks at the 3’ end may be due to the untranslated region of the ortholog of *Mipp2* (LOC6551318), the nesting gene. The lack of RNA-Seq data also affects the Splice Junction Predictions track, so we did not find a splice junction for the region between the first and second exon, and the splice junction score for the second splice junction (JUNC00005563) was low (a score of 15). However, the *Drosophila* Conservation of 28 species track in *D. melanogaster* indicates high identity in the coding sequence between *D. melanogaster* and *D. erecta* (Gold box I, Figure 1D). Long-read RNA sequence data is required to confirm the current gene model of *raptor* in *D. erecta*.

## Methods

“Detailed methods including algorithms, database versions, and citations for the complete annotation process can be found in Rele et al. (2023). Briefly, students use the GEP instance of the UCSC Genome Browser v.435 (https://gander.wustl.edu; Kent et al., 2002; Navarro Gonzalez et al., 2021) to examine the genomic neighborhood of their reference IIS gene in the *D. melanogaster* genome assembly (Aug. 2014; BDGP Release 6 + ISO1 MT/dm6). Students then retrieve the protein sequence for the *D. melanogaster* reference gene for a given isoform and run it using *tblastn* against their target *Drosophila* species genome assembly on the NCBI BLAST server (https://blast.ncbi.nlm.nih.gov/Blast.cgi; Altschul et al., 1990) to identify potential orthologs. To validate the potential ortholog, students compare the local genomic neighborhood of their potential ortholog with the genomic neighborhood of their reference gene in *D. melanogaster*. This local synteny analysis includes at minimum the two upstream and downstream genes relative to their putative ortholog. They also explore other sets of genomic evidence using multiple alignment tracks in the Genome Browser, including BLAT alignments of RefSeq Genes, Spaln alignment of *D. melanogaster* proteins, multiple gene prediction tracks (e.g., GeMoMa, Geneid, Augustus), and modENCODE RNA-Seq from the target species. Detailed explanation of how these lines of genomic evidenced are leveraged by students in gene model development are described in Rele et al. (2023). Genomic structure information (e.g., CDSs, intron-exon number and boundaries, number of isoforms) for the *D. melanogaster* reference gene is retrieved through the Gene Record Finder (https://gander.wustl.edu/~wilson/dmelgenerecord/index.html; Rele et al., 2023).

Approximate splice sites within the target gene are determined using *tblastn* using the CDSs from the *D. melanogaste*r reference gene. Coordinates of CDSs are then refined by examining aligned modENCODE RNA-Seq data, and by applying paradigms of molecular biology such as identifying canonical splice site sequences and ensuring the maintenance of an open reading frame across hypothesized splice sites. Students then confirm the biological validity of their target gene model using the Gene Model Checker (https://gander.wustl.edu/~wilson/dmelgenerecord/index.html; Rele et al., 2023), which compares the structure and translated sequence from their hypothesized target gene model against the *D. melanogaster* reference gene model. At least two independent models for a gene are generated by students under mentorship of their faculty course instructors. Those models are then reconciled by a third independent researcher mentored by the project leaders to produce the final model. Note: comparison of 5’ and 3’ UTR sequence information is not included in this GEP CURE protocol.” (Gruys et al., 2025)

## Supporting information

Gene model data files

## Supplemental Files

1. Zip file containing a FASTA, PEP, GFF files for the gene model
2. Figure 1 in high resolution

## Metadata

Bioinformatics, Genomics, *Drosophila*, Genotype Data, New Finding

## Acknowledgements

We would like to thank Wilson Leung for developing and maintaining the technological infrastructure that was used to create this gene model and Laura K. Reed for overseeing the project. Thank you to FlyBase for providing the definitive database for *Drosophila melanogaster* gene models. Further, we would like to thank the editors and developers at the journal *microPublication: Biology* for assistance in developing the template for these single gene ortholog publications.

## References

Altschul SF, Gish W, Miller W, Myers EW, Lipman DJ. 1990. Basic local alignment search tool. J Mol Biol 215(3): 403–410. PMID: 2231712

Backlund AE, Laskowski LF, Trosdal E, Wittke-Thompson J, Yowler B, Rele CP. 2025. Gene model for the ortholog of raptor in Drosophila busckii. Manuscript submitted.

Bock, IR. and Wheeler, MR. (1972). The Drosophila melanogaster species group. Univ Texas Publs Stud Genet 7(7213): 1–102.

Chirn GW, Rahman R, Sytnikova YA, Matts JA, Zeng M, Gerlach D, Yu M, Berger B, Naramura M, Kile BT, Lau NC. 2015. Conserved piRNA Expression from a Distinct Set of piRNA Cluster Loci in Eutherian Mammals. PLoS Genet 11(11): e1005652. PMID: 26588211

Gramates LS, Agapite J, Attrill H, Calvi BR, Crosby M, dos Santos G Goodman JL, Goutte-Gattat D, Jenkins V, Kaufman T, Larkin A, Matthews B, Millburn G, Strelets VB, and the FlyBase Consortium. 2022. FlyBase: a guided tour of highlighted features. Genetics 220(4): iyac035. PMID: 35266522

Grewal SS. Insulin/TOR signaling in growth and homeostasis: a view from the fly world. 2009. Int J Biochem Cell Biol 41(5):1006–1010. PMID: 18992839

Gruys ML, Sharp MA, Lill Z, Xiong C, Hark AT, Youngblom JJ, Rele CP, Reed LK. 2025. Gene model for the ortholog of Glys in Drosophila simulans. microPubl Biol:10.17912/micropub.biology.001168. PMID: 39845267

Hatfield I, Harvey I, Yates ER, Redd JR, Reiter LT, Bridges D. 2015. The role of TORC1 in muscle development in Drosophila. Sci Rep 5: 9676. PMID: 25866192.

Hietakangas V, Cohen SM. Regulation of tissue growth through nutrient sensing. 2009. Annu Rev Genet 43:389–410. PMID: 19694515

Jenkins VK, Larkin A, Thurmond J, FlyBase Consortium. 2022. Using FlyBase: A Database of Drosophila Genes and Genetics. Methods Mol Biol 2540: 1–34. PMID: 35980571

Kent WJ, Sugnet CW, Furey TS, Roskin KM, Pringle TH, Zahler AM, Haussler D. 2002. The Human Genome Browser at UCSC. Genome Res 12: 996–1006. PMID: 12045153

Larkin A, Marygold SJ, Antonazzo G, Attrill H, dos Santos G, Garapati PV, Goodman JL, Gramates LS, Millburn G, Strelets VB, Tabone CJ, and Thurmond J and the FlyBase Consortium. 2021. FlyBase: updates to the Drosophila melanogaster knowledge base. Nucleic Acids Res 49(D1): D899–D907. PMID: 33219682

Lee G, Chung J. 2007. Discrete functions of rictor and raptor in cell growth regulation in Drosophila. Biochem Biophys Res Commun 357(4): 1154–1159. PMID: 17462592

Lieser BC, Lose B, Kiser CA, Laskowski LF, Larsen CIS, Huber R, Thompson JS, Arsham AM, Chandrasekaran V, Rele CP. 2024. Gene model for the ortholog of Ilp4 in Drosophila erecta. Submitted.

Ma S, Avanesov AS, Porter E, Lee BC, Mariotti M, Zemskaya N, Guigo R, Moskalev AA, Gladyshev VN. 2018. Comparative transcriptomics across 14 Drosophila species reveals signatures of longevity. Aging Cell 17(4):e12740. PMID: 29671950

Markow TA and O’Grady P. 2005. Drosophila: A guide to species identification and use. London: Academic Press. ISBN: 978-0-12-473052-6

Mudge JM, Harrow J. 2016. The state of play in higher eukaryote gene annotation. Nat Rev Genet 17: 758–772. PMID: 27773922

Myers A, Hoffman A, Natysin M, Arsham AM, Stamm J, Thompson JS, Rele CP, Reed LK. 2024. Gene model for the ortholog Myc in Drosophila ananassae. microPubl Biol:10.17912/micropub.biology.000856. PMID: 39677519

Navarro Gonzalez J, Zweig AS, Speir ML, Schmelter D, Rosenbloom KR, Raney BJ, Powell CC, Nassar LR, Maulding ND, Lee CM et al. 2021. The UCSC Genome Browser database: 2021 update. Nucleic Acids Res 49(1): 1046–1057. PMID: 33221922

Raney BJ, Dreszer TR, Barber GP, Clawson H, Fujita PA, Wang T, Nguyen N, Paten B, Zweig AS, Karolchik D, Kent WJ. 2014. Track data hubs enable visualization of user-defined genome-wide annotations on the UCSC Genome Browser. Bioinformatics 30(7):1003–1005. PMID: 24227676

Rele CP, Sandlin KM, Leung W, Reed LK. 2023. Manual annotation of Drosophila genes: a Genomics Education Partnership protocol. F1000Research 11: 1579. 10.12688/f1000research.126839.2

Sturtevant AH. On the Subdivision of the Genus Drosophila. 1939. Proc Natl Acad Sci U S A 25(3):137–141. PMID: 16577879

Tello-Ruiz MK, Marco CF, Hsu FM, Khangura RS, Qiao P, Sapkota S, Stitzer MC, Wasikowski R, Wu H, Zhan J et al. 2019. Double triage to identify poorly annotated genes in maize: The missing link in community curation. PLoS One 14: e0224086– e0224013. PMID: 31658277

Tsacas L and Lachaise D. 1974. Les Drosophilidae des savanes preforestieres de la region tropicale de Lamto (Cote-d’Ivoire). II. Le peuplement des fruits de Pandanus candelabrum (Pandanacees). Annls Univ Abidjan E Ecol 7: 153–192.

Unwin AH. 1920. West African forests and forestry. E.P. Dutton & Co. Open Library Identifier: OL17442447M.

Wang T, Blumhagen R, Lao U, Kuo Y, Edgar BA. 2012. LST8 Regulates Cell Growth via Target-of-Rapamycin Complex 2 (TORC2). Mol Cell Biol 32(12): 2203–2213. PMID: 22493059

